# *CorShrink*: Empirical Bayes shrinkage estimation of correlations, with applications

**DOI:** 10.1101/368316

**Authors:** Kushal K. Dey, Matthew Stephens

## Abstract

Estimation of correlation matrices and correlations among variables is a ubiquitous problem in statistics. In many cases – especially when the number of observations is small relative to the number of variables – some kind of shrinkage or regularization is necessary to improve estimation accuracy. Here, we propose an Empirical Bayes shrinkage approach, `CorShrink`, which adaptively learns how much to shrink correlations by combining information across all pairs of variables. One key feature of `CorShrink`, which distinguishes it from most existing methods, is its flexibility in dealing with missing data. Indeed, `CorShrink` explicitly accounts for varying amounts of missingness among pairs of variables. Numerical studies suggest `CorShrink` is competitive with other popular correlation shrinkage methods, even when there is no missing data. We illustrate `CorShrink` on gene expression data from GTEx project, which suffers from extensive missing observations, and where existing methods struggle. We also illustrate its flexibility by applying it to estimate cosine similarities between word vectors from *word2vec* models, thereby generating more accurate word similarity rankings.

## 1 Introduction

Estimating the correlation matrix of a set of variables is a fundamental problem in statistics. The simplest estimator, the sample correlation matrix, can perform poorly when the number of variables (*p*) is large compared with the number of samples (*n*). This problem has motivated many alternative estimators, most of them based on shrinkage or regularization methods. These include using a convex combination of the sample correlation matrix with one or more target correlation matrices [Lancewicki and Aladjem, 2014, Ledoit and Wolf, 2003, 2004, Schafer and Strimmer, 2005, Touloumis, 2015]; thresholding-based approaches using either hard-thresholding [Bickel and Levina, 2008] or soft-thresholding [Rothman *and others*, 2009]; and L1-penalized shrinkage on correlation or inverse correlation matrices [Bien and Tibshirani, 2011, Friedman and *others*, 2008]. These methods, though often a considerable improvement on the sample correlation matrix, have their own limitations. In particular, almost all of these approaches assume that there are no missing observations. Here we present `CorShrink`, a fast and simple shrinkage-based approach to estimating correlations that can deal with missing observations, and, in many settings, is competitive in accuracy with existing methods.

In brief, `CorShrink` is a model-based extension of the approaches of Bickel and Levina [2008] and Rothman *and others* [2009]. Those approaches perform shrinkage using simple thresholding rules applied independently to each element of the correlation matrix. Our approach also treats each element of the matrix as independent, but uses an Empirical Bayes (EB) shrinkage approach that combines information across all elements to learn *how much to shrink.* Within this EB approach, the amount of shrinkage of each matrix element depends on the curvature of its likelihood, and so it naturally adapts to differing amounts of data available on each element. Our methods exploit semi-parametric EB methods [Stephens, 2016] that are both computationally fast and stable, and avoid strong assumptions on the distributions of pairwise correlations.

We illustrate `CorShrink` on a dataset with extensive missingness: gene expression data across 53 human tissues (and cell-lines) collected by Genotype Tissue Expression (GTEx) project [Lonsdale *and others*, 2013]. The data matrix for each gene consists of expression measurements on many post-mortem donors across many tissues. However, for most donors, data are available on only a subset of tissues, and many elements of this data matrix are missing. Some tissues are particularly rarely sampled, and so their sample correlations are particularly noisy. Most existing correlation matrix estimation procedures cannot be easily applied to these data. In contrast, as we illustrate here, `CorShrink` is straightforward to apply. Further, the resulting shrinkage estimates are more biologically plausible, and visually considerably less cluttered, than the sample correlation matrix estimate. We also compare results with an alternative approach based on first imputing the missing values in the data matrix, and conclude that for data like these with high levels of missingness this imputation strategy is unappealing.

In addition to these results with missing data, we also use simple simulations to show that even when no data are missing, `CorShrink` is often competitive in accuracy with other methods, especially when the true correlation matrix is sparse. Furthermore, `CorShrink` is easy to apply to other correlation-like quantities, and we illustrate this by applying it to cosine similarities of vectors from *word2vec* models [Mikolov *and others*, 2013] that measure word-word similarities.

## 2 Methods

Let (*X_np_*)_*N*×*p*_ denote a data matrix with *N* samples and *P* variables, where some values may be missing (recorded as NA). For each pair of variables *i, j* ∈ {1, 2, ···, *P*} let *R_ij_* denote their (unknown) true correlation, and 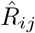 denote the sample correlation computed using only the samples *n* that have observed values for both the variables *i* and *j* (e.g. using the option `use=“pairwise.complete.obs”, method = “pearson”` in the R function `cor`). Further, let *Z_ij_* and *Ẑ_ij_* denote the corresponding Fisher Z-transforms [Fisher, 1915]:

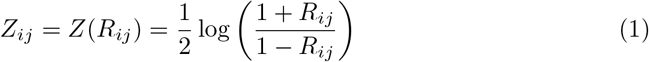

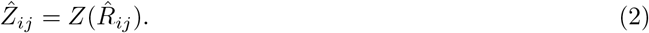

Our goal is to estimate the matrix *Z* = (*Z_ij_*) and hence *R* = (*R_ij_*) from the data *Ẑ* = (*Ẑ_ij_*).

Our starting point is a result from Fisher [1921], who showed that, under a bivariate normality assumption on each pair of variables, the observations *Ẑ_ij_* are approximately normal:

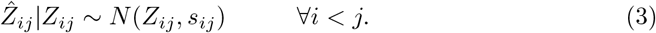

Here the standard deviation *s_ij_* is given by

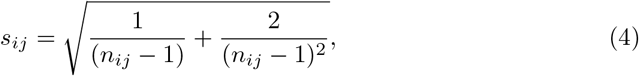

where *n_ij_* > 3 is the number of *matched samples*, for which both variables *i* and *j* are observed:

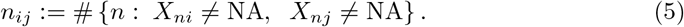

We take an Empirical Bayes approach, combining the likelihood (3) with an assumption that the *Z_ij_* are drawn from some underlying distribution *g*:

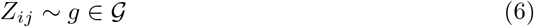

where 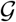 denotes some pre-specified family of distributions. There are many possible choices for 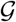, and here we make use of the flexible nonparametric methods implemented in the R package `ashr` [Stephens, 2016]. These include:

1. 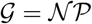, the set of all real-valued distributions. This is the fully non-parametric approach from Koenker and Mizera [2014];
2. 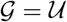 the set of all unimodal distributions;
3. 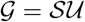 the set of all symmetric unimodal distributions;
4. 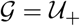 (respectively 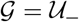) the set of unimodal distributions that are constrained to be positive (respectively negative).
5. 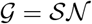 the set of all scale-mixtures of normals.

The rationale for the more restrictive unimodal choices 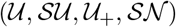 is that they ensure shrinkage towards a common point, and also regularize the estimate of *g* compared with a fully non-parametric approach. Stephens [2016] focused on the case where the mode of *g* was at 0, but the `ashr` package also implements methods to estimate the location of the mode, which may also be useful in this context. Results in this paper were generated using 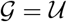 with mode at 0 unless otherwise stated.

We now make two simplifying assumptions: that the *Z_ij_* are independent from (6) and *Ẑ_ij_*|*Z_ij_* are independent from (3). Neither of these assumptions is true, but composite likelihood theory [Varin *and others*, 2011] suggests that point estimates of both *g* and *Z* should be somewhat robust to dependence, and our numeric experiments later show that point estimates from our method can outperform methods that do model the dependence. These simplifying assumptions can be seen as a model-based analogue of previous methods that apply simple thresholding rules independently to each element of the correlation matrix [Bickel and Levina, 2008, Rothman *and others*, 2009].

We fit the above model to obtain the posterior mean of *Z_ij_*, 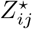, given 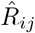:

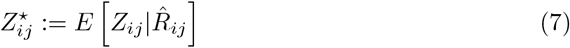

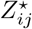 are adaptively shrunk estimates of *Ẑ_ij_* that account for *n_ij_*, the number of matched samples between variables *i* and *j*. The smaller the *n_ij_*, the larger would be *s_ij_* in Equation 4 and higher would be the level of shrinkage on *Ẑ_ij_*.

Next, we reverse transform 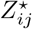 to correlation estimate 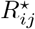:

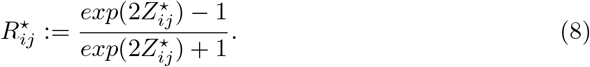

Just as with simple threholding methods the matrix 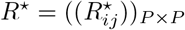 may not be positive definite, so we select the nearest positive definite matrix *R^⋆⋆^* to *R^⋆^*, using the method from Higham [2002].

If the variables are not pairwise normally distributed, then the representation of *s_ij_* as in Equation (4) does not hold. One approach in this context is to use transformations of the data that are more robust to the non-normality of the data, for example – Box-cox, ranks, rank-based inverse normal (RIN) transformations [Bishara and Hittner, 2015, 2017]. Another approach would be to estimate s_ij_· using Bootstrap methods [Diaconis and Efron, 1983, Efron, 1981]. The flexibility to use re-sampling methods as above, extends the scope of the `CorShrink` method beyond correlations to any correlation-like quantities-partial correlations, rank correlations, cosine similarities between word vectors in a word2vec model etc.

## 3 Results

### 3.1 Applications – Genetics

We first illustrate `CorShrink` on a data set with many missing observations. The Genotype Tissue Expression (GTEx) Project [Lonsdale *and others*, 2013] collected gene expression data from hundreds of post-mortem donors across many different tissues. The version of the data we consider here (v6) contains ~ 540 donors across 51 different tissues and 2 cell lines. If every donor contributed every tissue then the expression data at each gene would be a 540 by 53 matrix, each entry of which is a real number (a measure of expression known as log-counts-per-million; Law *and others* [2014]). Different donors, however, contributed different tissues leading to a large number of missing observations (> 70%) in this matrix.

Figure 1 illustrates `CorShrink` using data from a single gene, *PLIN1.* Panel (a) shows the pairwise sample correlation between tissues, which are input to `CorShrink`. One notable feature of these sample correlations is strong positive correlation among brain-derived tissues. Another feature is that many of the sample correlations are quite strongly negative. Biologically, strong negative correlation among tissues are expected to be rare, so these observations may be driven primarily by sampling variation. This view is further supported by noting that the most negative values tend to occur in tissue pairs that have fewer matching samples (e.g. nerve-tibial and cervix-endocervix have correlation −0.89 with only 3 matching samples). Panel (b) shows the estimated *ĝ*, with 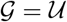 and the mode fixed to 0. The estimate is skewed towards small positive values, consistent with the biologically plausible scenario that most pairs of tissues are mildly positively correlated. Panel (c) shows the resulting `CorShrink` shrinkage estimates of the correlation. The sample correlations are all shrunk towards the mode of *g*, but the amount of shrinkage for each tissue pair depends on matched sample sizes: pairs with few matched samples undergo strong shrinkage while those with more matched samples remain largely unperturbed (Panel (d)). The negative sample correlations are mostly removed in the shrinkage estimates – due to both the concentration of *ĝ* on positive values, and the stronger shrinkage of tissue pairs with small matched sample size.

Given the biological expectation that negative correlations among tissues are unlikely, we might prefer to constrain the correlations to be positive here by using 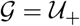. The results for this, in addition to several other choices of 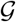 are shown in Supplementary Figure S1. The results in this case are largely consistent across these choices of 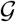, although only 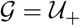 completely removes negative estimates.

These illustrative results were “gene-based” – that is, they were based on data at the *PLIN1* gene alone, and involved shrinking correlations for all tissue-pairs using the same prior distribution *ĝ*. However, for some tissue pairs (e.g. biologically-related tissues, like the two heart tissues) strong correlations are more plausible than for others. Here, since we have data on many genes for each tissue pair, we can instead separately estimate a prior distribution *ĝ* for each tissue pair, by combining data across genes. That is, for each tissue pair, we feed into Equation 3 the vector of pairwise correlation in expression for this tissue pair *across all genes*, together with their corresponding standard errors. Supplementary Figure S2 shows results from this “tissue-pair-based shrinkage” with the “gene-based shrinkage” in Figure 1. The two results are reassuringly similar, especially when compared with the original sample correlations.

**Figure 1.**
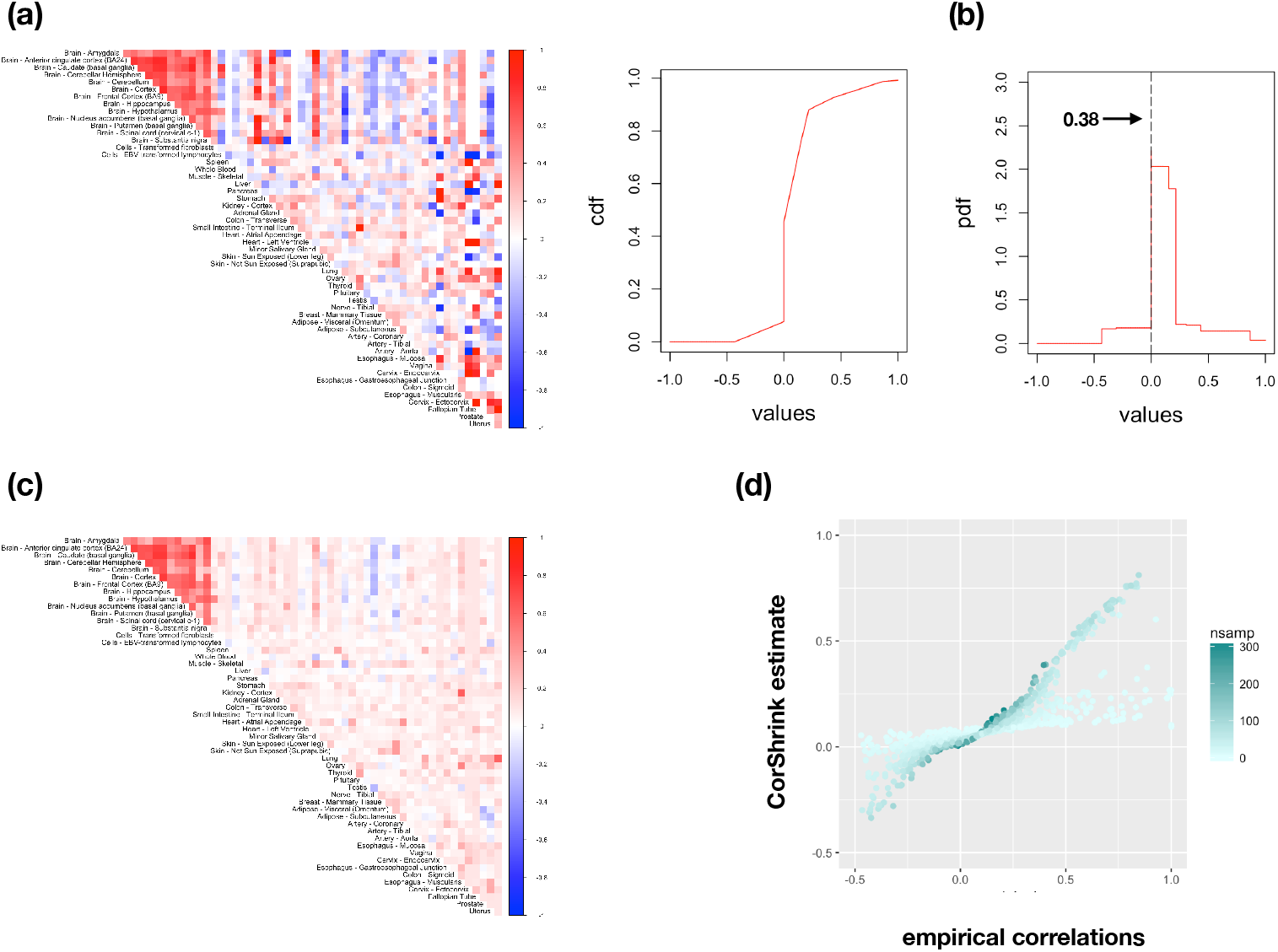
Illustration of `CorShrink` in action. (a) The image plot of the pairwise correlation matrix between tissue pairs for the log CPM expression data [Law *and others*, 2014] of the *PLIN1* gene. (b) The probability and cumulative density function plots for the EB shrinkage prior in `CorShrink` generated using 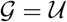. 0.38 is the empirically determined prior probability mass of observing 0 correlation. (c) The image plot of the estimated correlation matrix using `CorShrink`. The representation is visually more parsimonious and arguably more interpretable than (a). (d) Scatter plot of the pairwise sample correlation values against the `CorShrink` fitted estimates for all tissue pairs. Each point is colored by the number of postmortem donors contributing the corresponding tissue pair. Expectedly, the pairs with low numbers of matching samples (light colored points) undergo high shrinkage while those with larger number of matching samples remain largely unperturbed.

To further illustrate the flexibility of `CorShrink`, Figure 2 shows results for three different genes that show different qualitative patterns of correlation. One gene (*VSIR*) shows correlation patterns similar to those for *PLIN1* above, with strong correlations mostly among brain tissues; this type of pattern occurs in many genes in this data set (not shown). In contrast the *HBB* gene (which encodes beta-globin, a sub-unit of hemoglobin) shows high correlations across many tissue pairs – both brain, and non-brain. We noticed similar patterns in other hemoglobin-related genes, *HBA1* and *HBA2* (not shown). Finally, the gene *MTURN* shows little correlation in expression across most pairs of tissues. These results particularly highlight the adaptiveness of the gene-based analysis to gene-specific patterns. Effectively `CorShrink` learns that the sample correlations in *MTURN* are mostly consistent with weak correlation, and so shrinks these sample correlations strongly toward 0. By comparison the shrinkage in *HBB* is considerably less pronounced.

**Figure 2.**
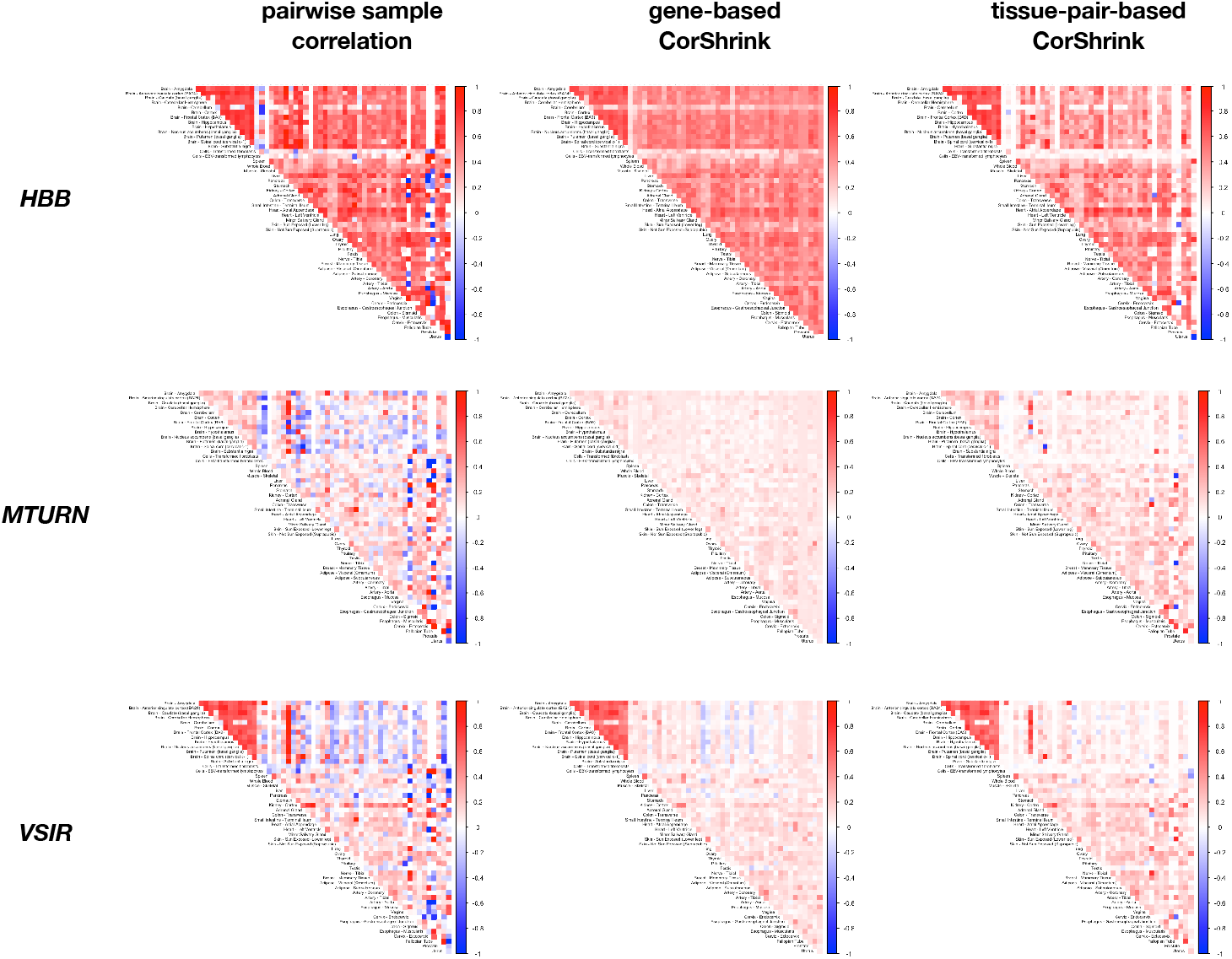
Results from `CorShrink` applied to three different genes (*HBB, MTURN, VSIR*) showing qualitatively different patterns of correlation. The gene-based results illustrate how `CorShrink` adapts itself to different amounts of correlation in different genes (e.g. compare the modest shrinkage for *HBB* with the stronger shrinkage for *MTURN*). See text for further discussion.

One important feature of `CorShrink` is its ability to deal with a high rate of missing data (> 70% in these examples). One general and widely-used approach to deal with missing data is to impute them, for example by using low-rank-based methods such as softImpute [Hastie and Mazumder, 2015, Mazumder *and others*, 2010] or FLASH [Wang and Stephens, 2018]. However, we have found that in this context, with its high rate of missing data, imputation can grossly distort correlation estimates. For example, Figure 3 illustrates the dramatic effects of imputation for *PLIN1*. The imputed data from both softImpute and FLASH show a strong upward bias in their correlation for most tissue pairs, which appears inconsistent with the observed pairwise correlations.

**Figure 3.**
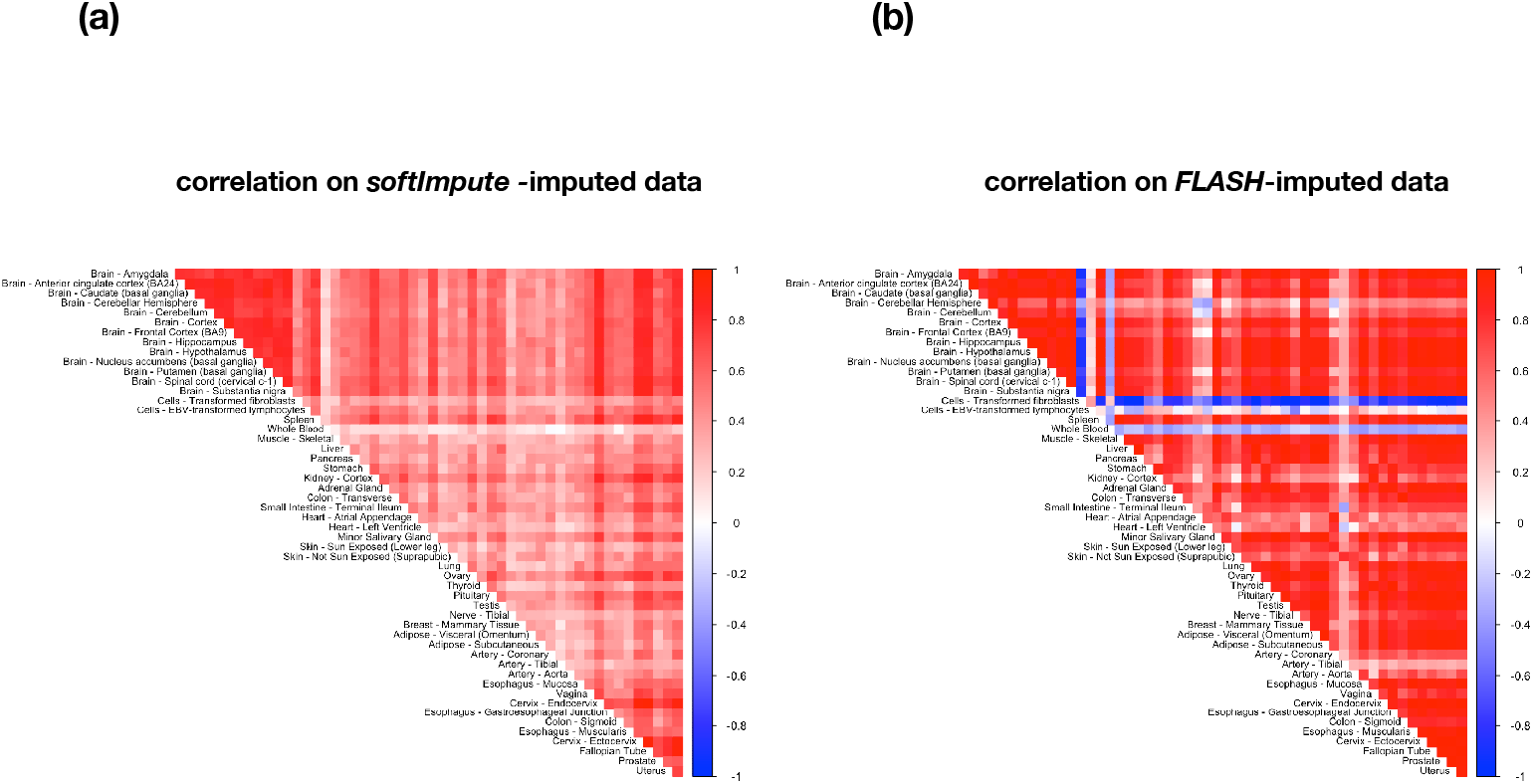
Illustration of impact of imputation on correlation matrix estimates. The panels show sample correlation matrices for the same data (*PLIN1*) as in Figure 1, after imputation using (a) softImpute [Mazumder *and others*, 2010] and (b) FLASH Wang and Stephens [2018]. In both cases the results appear inconsistent with the observed data, unlike the results from `CorShrink` (Figure 1a, c)

### 3.2 Simulation studies

Although `CorShrink` is particularly well-suited to settings with missing data, it is also a very competent correlation shrinkage method for complete data. Here we demonstrate this using simulations to compare `CorShrink` with other correlation shrinkage methods: soft thresholding estimator *PDSCE* [Rothman, 2012] and *corpcor*, which performs shrinkage towards the identity matrix [Schäfer and Strimmer, 2004, 2005]. We also compare with *GLASSO* [Friedman *and others*, 2008, Meinshausen and Buhlmann, 2006, Witten *and others*, 2011], which is based on inducing sparsity in the precision matrix.

We simulated multivariate normal data with 0 mean and three types of correlation structure: a hub correlation matrix (sparse correlation, sparse precision), a Toeplitz correlation matrix (sparse correlation, non-sparse precision) and a banded precision matrix (non-sparse correlation, sparse precision). In each case we simulated *p* = 100 features for sample size *n* = 30, 50,100,1000. We ran `CorShrink` with 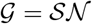; we ran *GLASSO* at varying levels of the tuning parameter λ = 0.01,0.1, 0.5 and 1.

Figure 4 compares the accuracy of each method in estimating the correlation matrix, as measured by Correlation Matrix Distance (CMD) [Herdin *and others*, 2005] between the true and estimated matrix. For both the hub and Toeplitz scenarios `CorShrink` was consistently the most accurate, and for the sparse banded precision matrix case it performed similarly to the best-performing of the *GLASSO* approaches. Similar results hold if accuracy is measured using Frobenius distance instead of CMD (Supplementary Figure S3).

**Figure 4.**
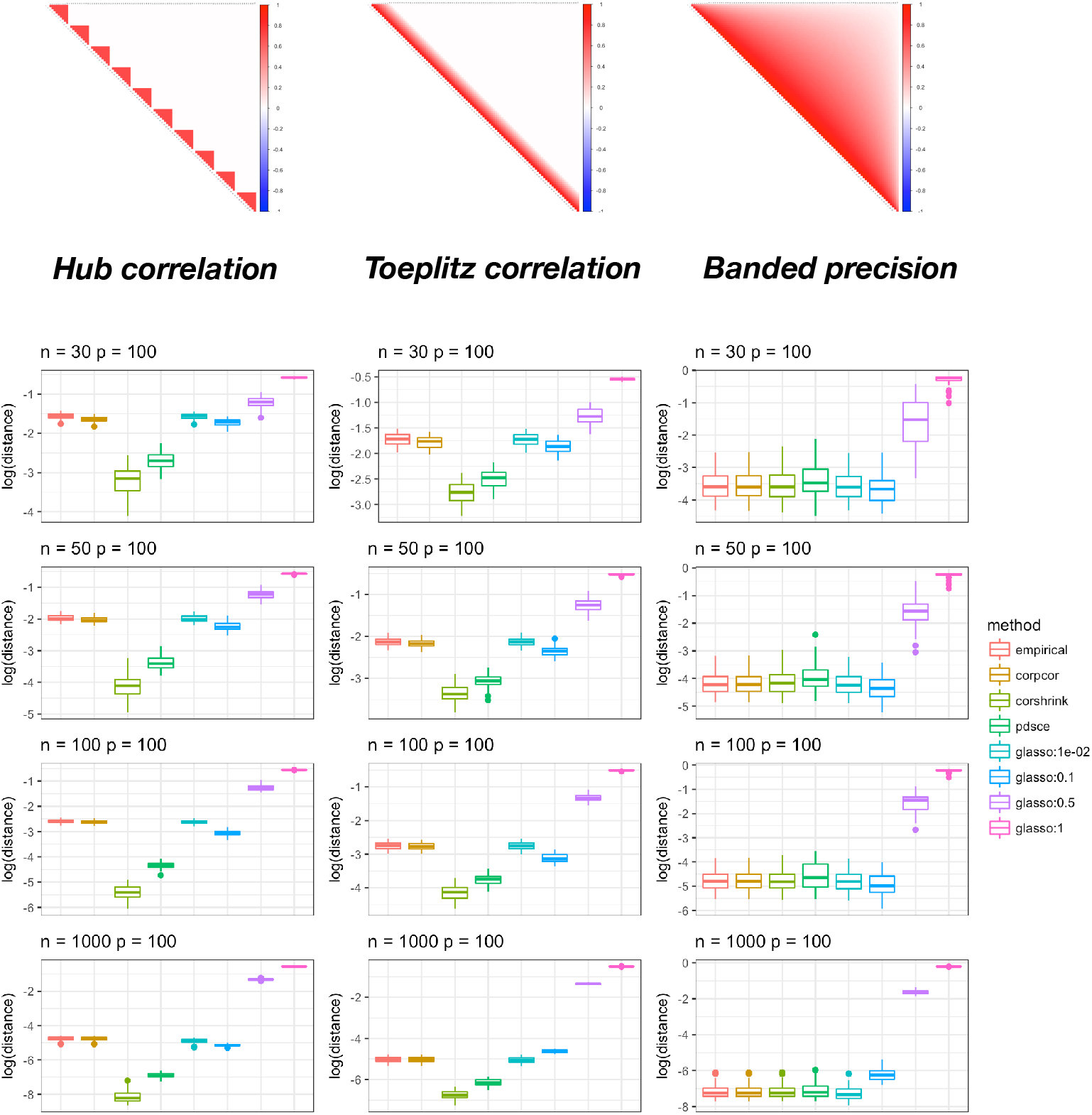
Box plot of the Correlation Matrix Distance (CMD) [Herdin *and others*, 2005] between population correlation matrix and the estimated matrix from different methods *-corpcor*[Schafer and Strimmer, 2004, 2005], `CorShrink`, *PDSCE* [Rothman, 2012], GLASSO [Friedman *and others*, 2008] and the empirical pairwise correlation matrix. Three types of population correlation structure were considered – hub structure, Toeplitz structure and the banded precision matrix structure. Image plots for these population correlation matrices are shown in the topmost panel. GLASSO was fitted using a broad range of tuning parameters (λ = 0.01, 0.1, 0.5 and 1), and their relative performance was also compared. `CorShrink` outperforms the other methods for the sparse structured population correlation model examples (hub and Toeplitz), closely followed by *PDSCE.*

Figure 5 compares, for each method, the largest eigenvalues of the estimated correlation matrices with those of the population correlation matrix. Again, for hub and Toeplitz scenarios, `CorShrink` estimates consistently follow the population eigenvalues more closely than other methods.

**Figure 5.**
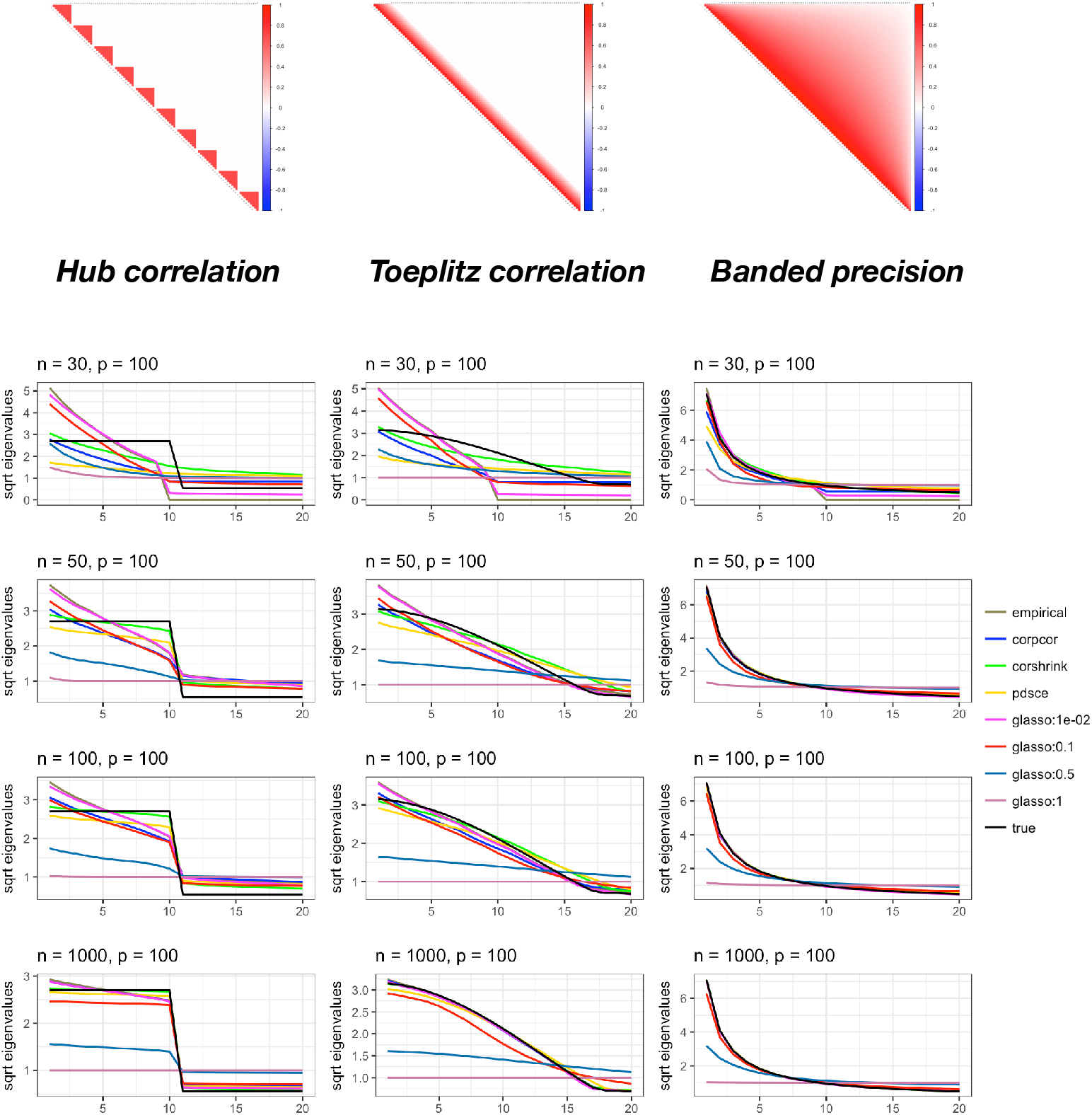
Plots of sorted square-root eigenvalue trends of the population correlation matrix against those of the estimated correlation matrices using different methods – *corpcor* [Schafer and Strimmer, 2004, 2005], `CorShrink`, *PDSCE* [Rothman, 2012], GLASSO [Friedman *and others*, 2008] and the empirical pairwise correlation matrix. Three types of population correlation structure were considered – hub structure, Toeplitz structure and the banded precision matrix structure. Image plots for these population correlation matrices are shown in the topmost panel. GLASSO was fitted using a broad range of tuning parameters (λ = 0.01, 0.1, 0.5 and 1), and their relative performance was also compared. The trend of sorted eigenvalues from `CorShrink` more closely follow the sorted population eigenvalues, in particular for the hub and the Toeplitz model examples.

In some applications interest may be focused on estimating the inverse of the correlation matrix, or the partial correlation matrix, rather than the correlation matrix itself. As currently implemented `CorShrink` is not well suited to such applications: in particular the correlation matrix estimate from `CorShrink` is typically close to singular when *n* << *p*, so its inverse is unstable, and does not generally provide a good estimate for the inverse-correlation matrix (in Frobenius norm, say). In contrast *GLASSO* is specifically designed for estimating (sparse) inverse correlation matrices, and indeed for this task *GLASSO* outperforms the other methods considered here on our examples (Supplementary Figure S4), which all involve sparse precision matrices.

### 3.3 Applications – Natural Language Processing

`CorShrink` can also be applied to perform shrinkage estimation of other correlation-like quantities. Here we illustrate this in an application from Natural Language Processing: estimating cosine similarities between vector representations of words, as obtained from a *word2vec* [Mikolov *and others*, 2013] or *GLOVE* [Pennington *and others*] model for example. The aim here is to generate more robust estimates of the cosine similarities between words, less affected by sampling variation due to some words appearing relatively infrequently in a corpus.

We obtained text data from the monthly issues of *Ebony* magazine published in 1968. This year marked a major moment in American history with the assassination of Dr. Martin Luther King (MLK) and the subsequent effect on the civil rights movement, and these events are reflected in many of the articles. We fitted *word2vec* on the combined text data from these issues, obtained vector representations and computed (sample) cosine similarities between words based on the vector representations. We transformed these to Fisher Z-scores using (1).

Unlike correlations, there is no obvious way to compute standard errors of the Fisher Z-scores for these cosine similarities. Consequently we used re-sampling methods – specifically 100 bootstrap samples [Diaconis and Efron, 1983, Efron, 1981], with magazine issues being the sampling unit – to estimate standard errors. We used issues rather than articles as the sampling unit to preserve the feature that each issue typically contains a range of different article types (e.g. on sports, politics, lifestyle etc).

We focused on three words/phrases: *martin luther king*, *civil rights* and *vietnam*, treating each as a single “word” for our analysis. For each of these phrases we applied `CorShrink` to the corresponding Fisher *Z*-scores, with re-sampling standard errors computed as above. Figure 6 shows the top words contextually similar to the focal phrases based on cosine similarities before and after `CorShrink` shrinkage.

**Figure 6.**
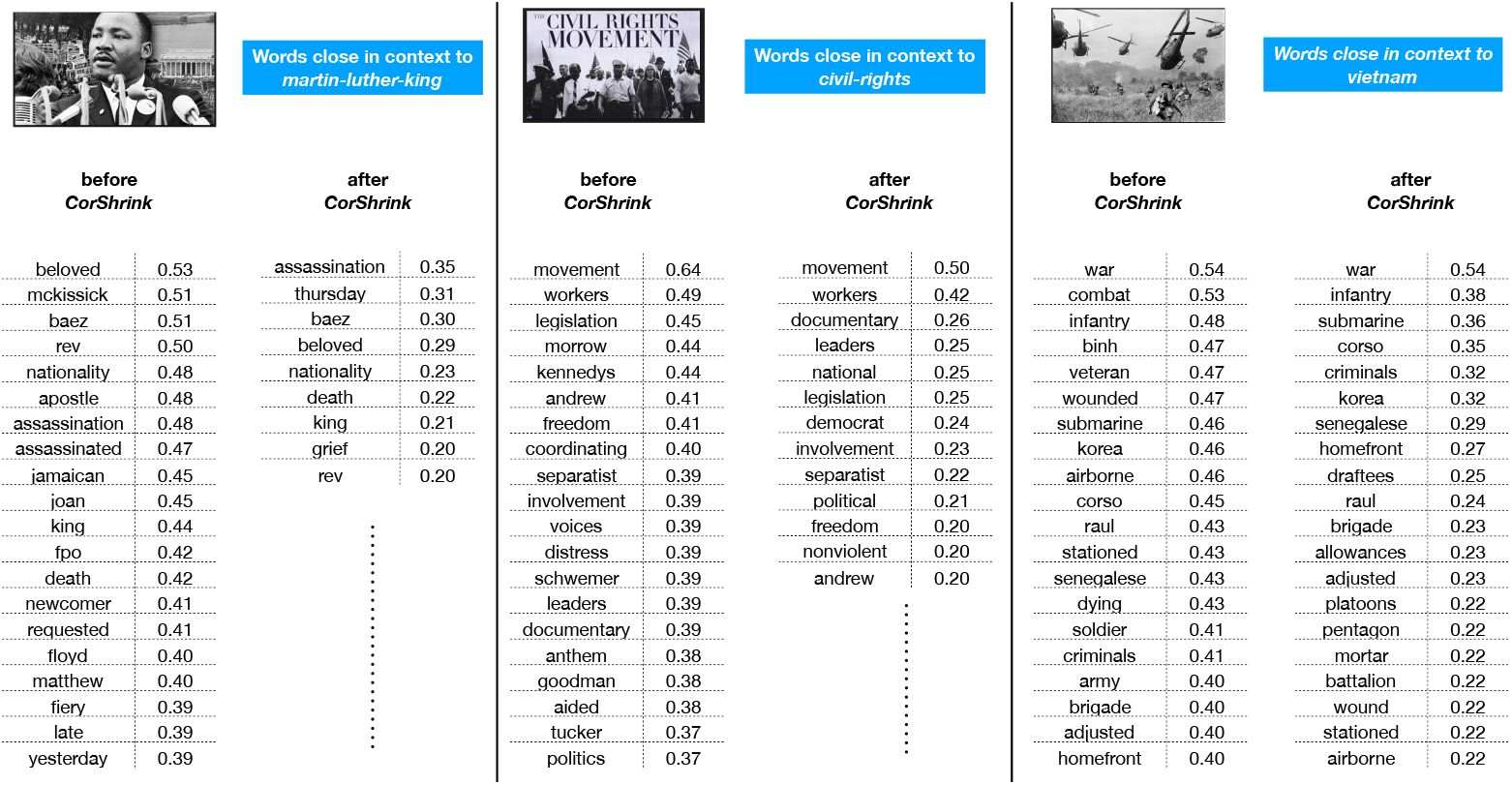
Results of `CorShrink` applied to estimates of cosine similarities between words. The tables show the top ≤ 20 words estimated to be most similar to the word/phrase (s) **martin-luther-king** (*left*), **civil-rights** (*middle*) and **vietnam** (*right*), before and after applying `CorShrink`. Only words with estimated similarity score > 0.2 are shown. The “before `CorShrink`” results were generated by running *word2vec* [Mikolov *and others*, 2013] on the monthly issues of the *Ebony* magazine in the year of 1968. Words/phrases that occur infrequently in the context of the focal word/phrase are shrunk more strongly by `CorShrink`.

As expected, the cosine similarities after `CorShrink` are consistently smaller, due to shrinkage towards 0. However, `CorShrink` also reorders the similarities because some words are shrunk more strongly than others. For example, the word *apostle*, which is strongly connected to the phrase *martin-luther-king* before `CorShrink` (ranked 6), has its cosine similarity shrunk strongly (from cosine similarity 0.48 to 0.13). This strong shrinkage occurs because the word occurred overall in 5 issues, and in only one of these issues was it very strongly associated with MLK (in the context of eulogizing him, immediately following his death in the May 1968 issue). In contrast, *Thursday*, which was the day of MLK’s assassination, was shrunk much less strongly (from cosine similarity 0.38 to 0.31), because it occurs overall in 2 issues and consistently in connection with MLK in the May 1968 issue.

This shrinkage helps to highlight the words that are truly very strongly associated. For example, in connection with MLK the word *assassination* appears much higher up the list post shrinkage (moving from 7th to 1st). And although the word *war* is very strongly associated with *vietnam* both before and after shrinkage, its connection is clearer post-shrinkage because other words like *combat* and *infantry* are shrunk more strongly.

## 4 Discussion

We presented `CorShrink`, an application of the adaptive shrinkage framework from [Stephens, 2016] to shrinkage estimation of correlation matrices and related quantities. Unlike most correlation matrix estimation approaches (*corpcor, GLASSO*), `CorShrink` can deal with missing values in the data matrix, and in this setting adjusts the degree of shrinkage based on the number of observations contributing to each matrix element (Figure 1). Even with no missing data, `CorShrink` can outperform the other methods when the true correlation matrix is sparse (see Figures 4, 5). In addition `CorShrink` has the flexibility to be applied to any set of correlation-like quantities, and not only correlation matrices. Here we illustrated this by shrinking cosine similarities between word pairs from text data (Figure 6).

The basic strategy underlying `CorShrink` is very simple, and involves shrinking each correlation term independently (after using all terms to learn how much to shrink). A similar simple strategy is also used in *corpcor*, and Bickel and Levina [2008] provide theoretical results for another (even simpler) strategy of this form. The simplicity of this strategy is very attractive, and is what allows `CorShrink` to easily deal with missing data for example. However, when the true correlation matrix is highly structured – for example, low rank plus diagonal – then methods that exploit this structure [Fan *and others*, 2016] may be more accurate.

In terms of speed, `CorShrink` is comparable to *corpcor*, and considerably faster than *glasso* and *PDSCE* when one takes account of the need to tune these methods using cross-validation. As noted in the Results, the current implementation of `CorShrink` is not suited to estimating inverse correlations, and it may be interesting to consider whether modifications to the approach could rectify this. `CorShrink` is available as an R package on CRAN and is also under active development on Github https://github.com/kkdey/CorShrink; code implementing the analyses presented here are available at https://kkdey.github.io/CorShrink-pages/.

---

**Figure S1.**
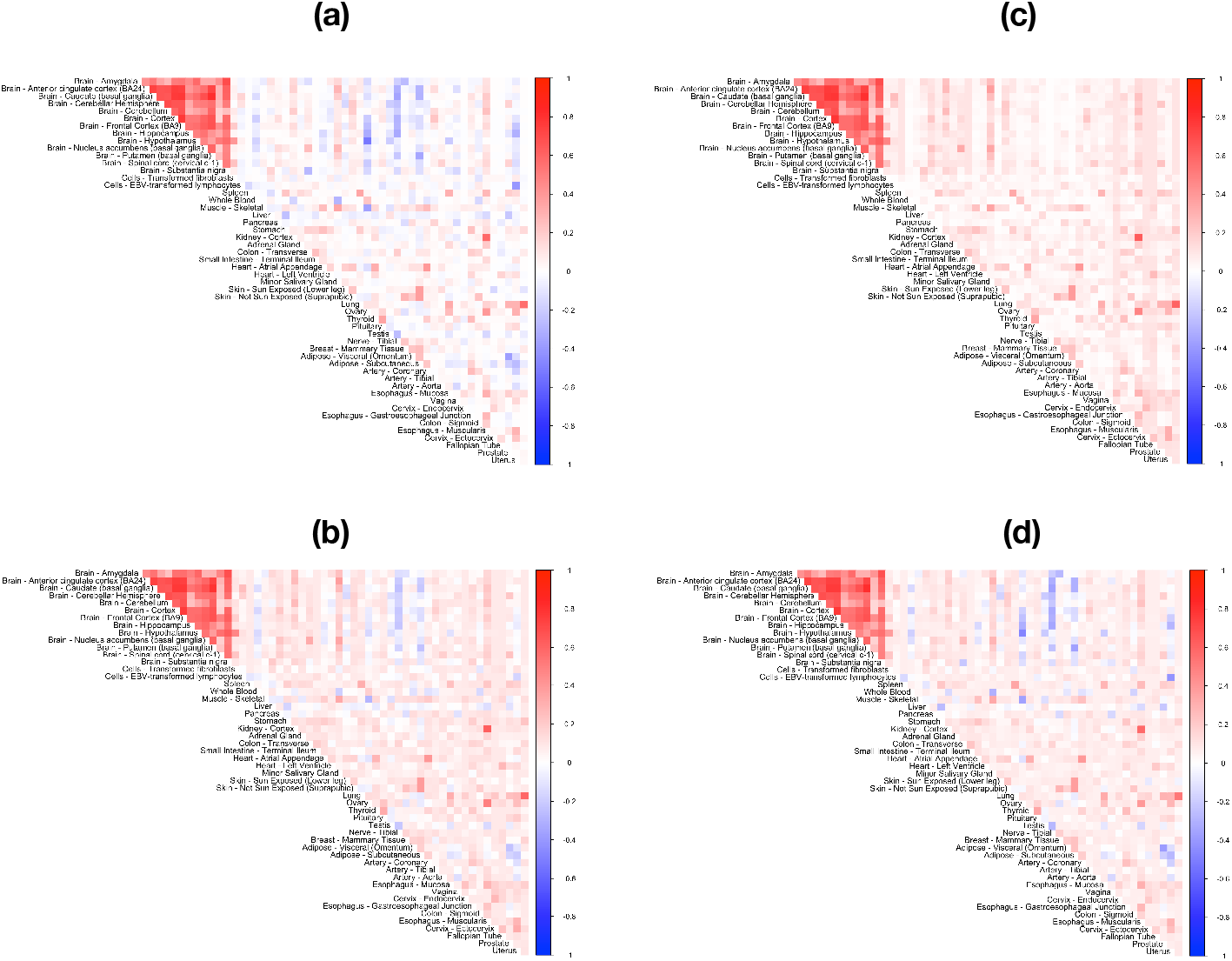
Image plots from applying `CorShrink` shrinkage on the donor by tissue expression data for *PLIN1* gene using choices of 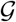 (in Equation 6) other than 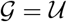 already reported in Fig 1. (a) 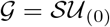 where 0 is the mode of the distribution. (b) 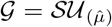 where the mode 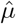 is not necessarily 0 and is estimated from the data. (c) 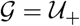 and (d) 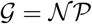. See text for explanation of terminology and discussion.

**Figure S2.**
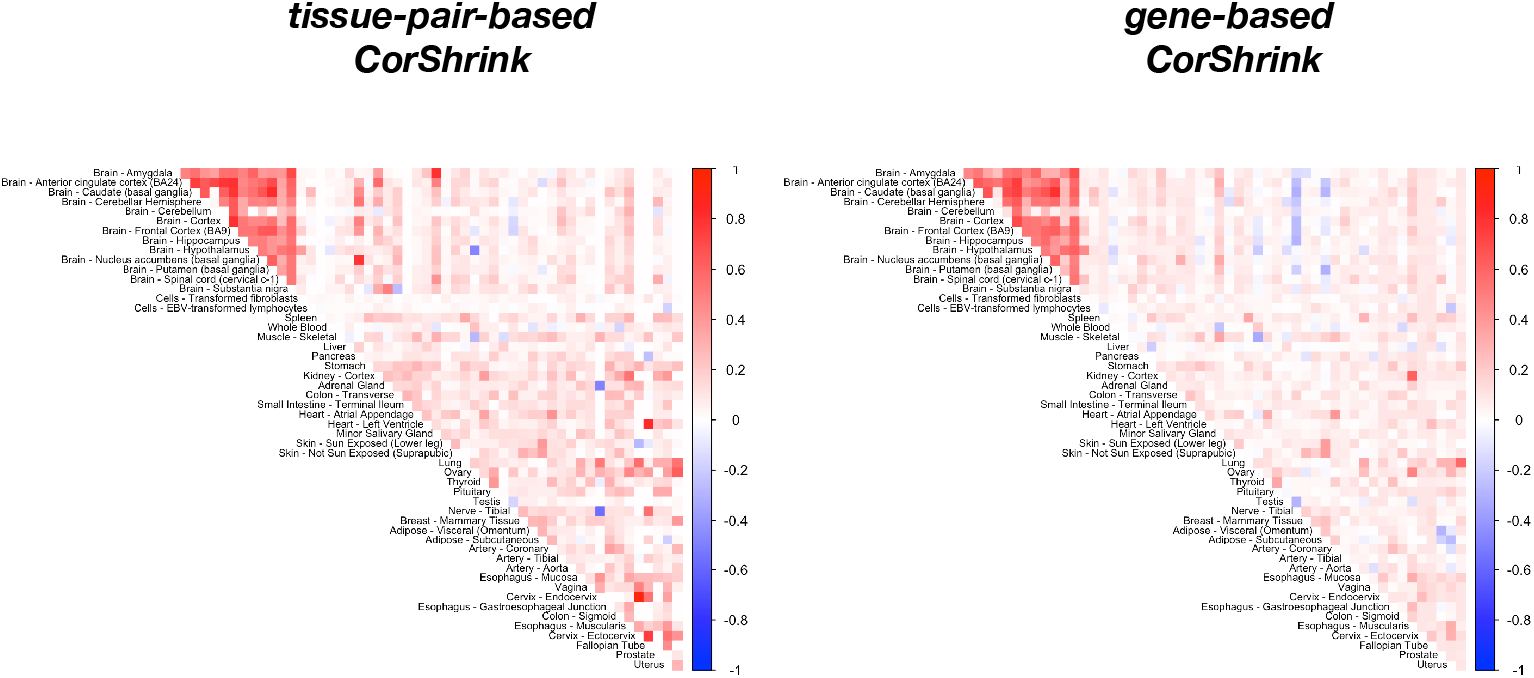
Image plots of the estimated correlation matrices after applying `CorShrink` on the subject by tissue expression matrix data for the *PLIN1* gene for two different models – (a) “‘tissue-pair-based shrinkage” and (b) “‘gene-based shrinkage” models and with 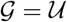. Both representations are re-assuringly similar.

**Figure S3.**
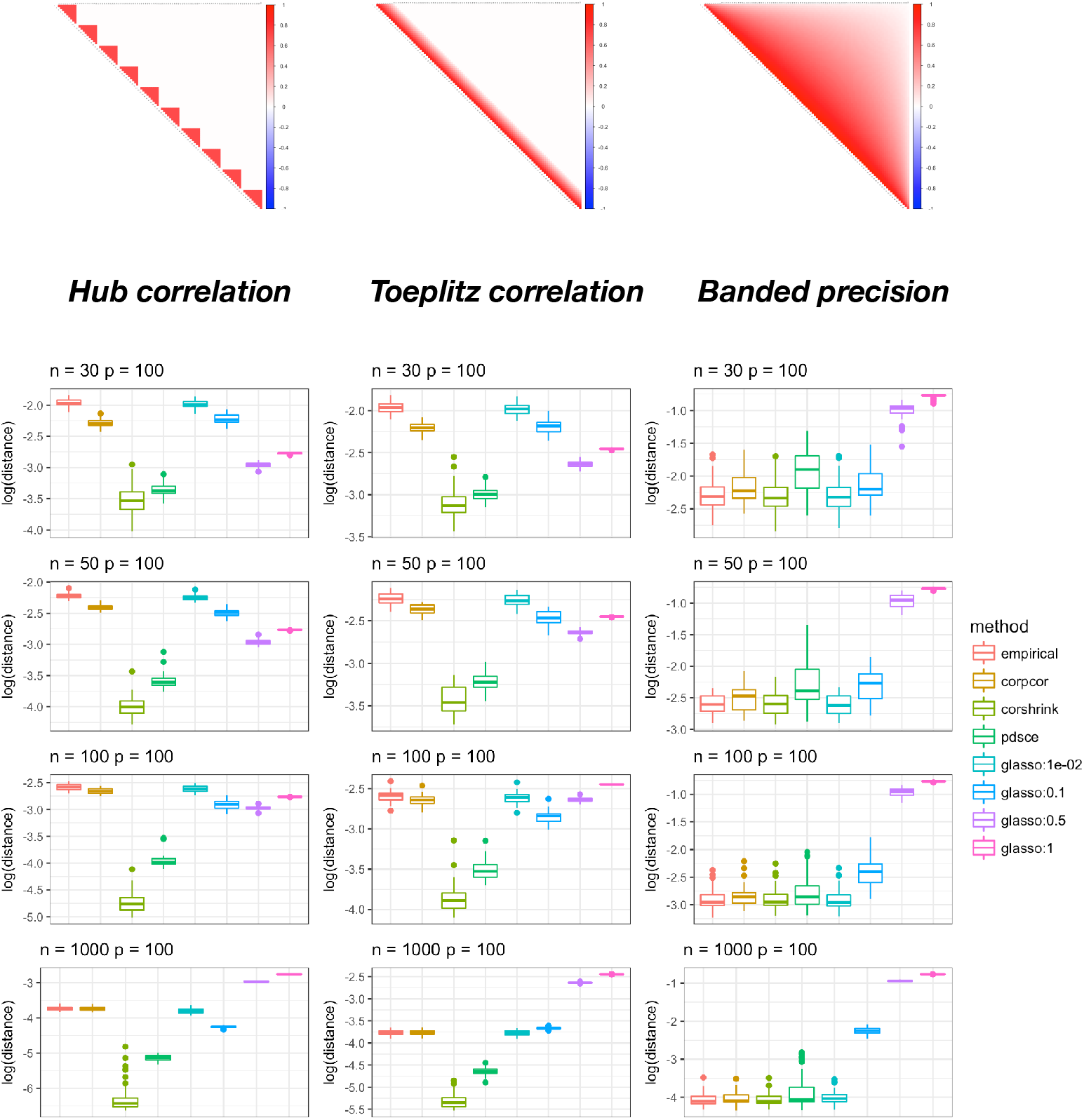
Box plot of the Frobenius distance between population correlation matrix and the estimated matrix using different methods – *corpcor* [Schäfer and Strimmer, 2004, 2005], `CorShrink`, *PDSCE* [Rothman, 2012], GLASSO [Friedman *and others*, 2008] and the empirical pairwise correlation matrix – for different structural assumptions on the underlying population correlation, hub structure, Toeplitz structure and the banded precision matrix structure. Image plots for the population correlation matrices are shown in the topmost panel. GLASSO was fitted using a broad range of tuning parameters, and their relative performance was also compared. `CorShrink` outperforms the other methods for the structured/sparse covariance models (hub and Toeplitz), closely followed by *PDSCE.*

**Figure S4.**
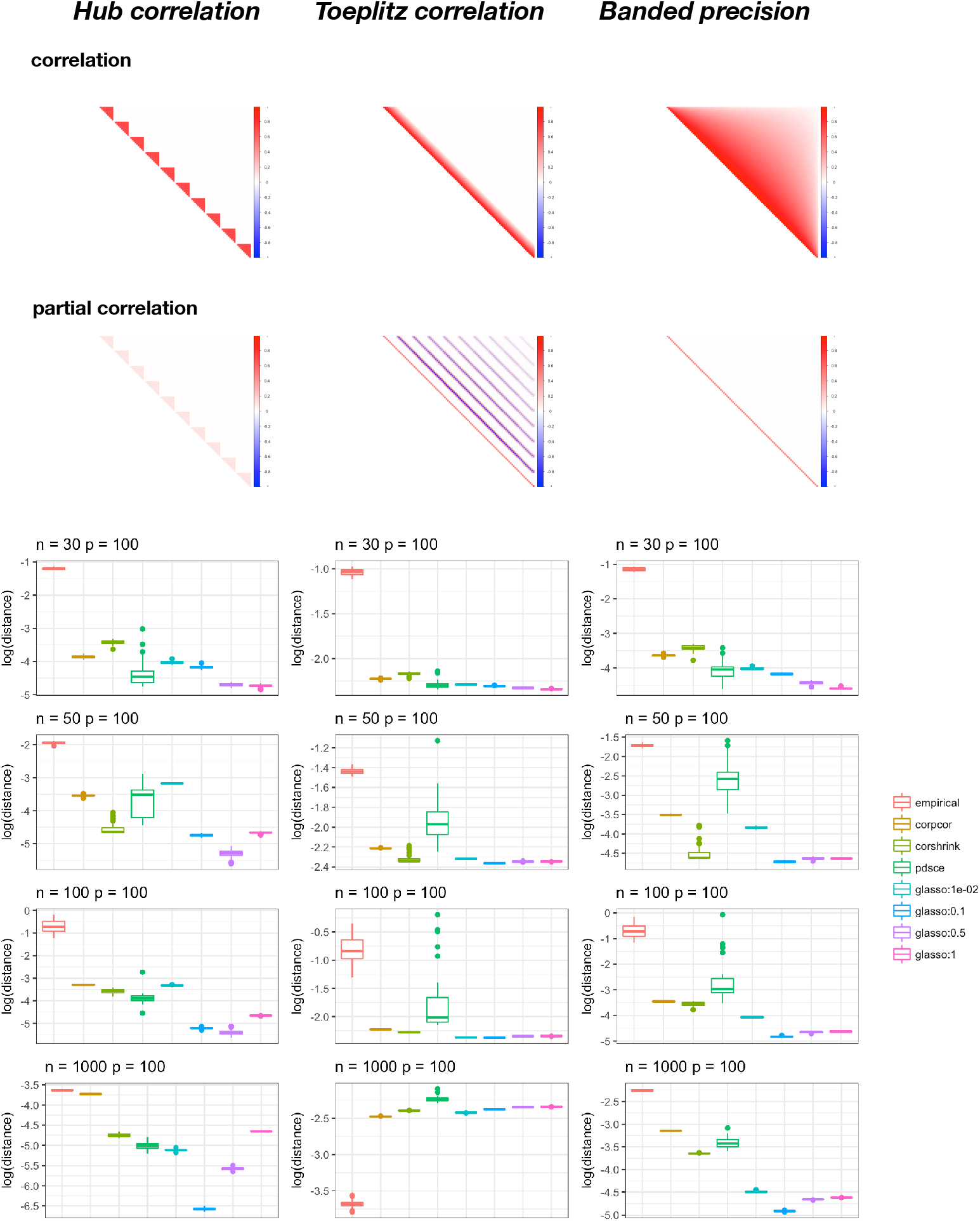
Comparing the accuracy (using Frobenius distance with respect to population partial correlation matrix) of the estimated partial correlation matrix from `CorShrink` relative to the partial correlation matrix obtained from correlation estimates of competing methods *-corpcor* [Schäfer and Strimmer, 2004, 2005], *PDSCE* [Rothman, 2012] and GLASSO [Friedman *and others*, 2008] – where the population correlation matrix profiles are as in Supplementary Figure S3 with image plots of these correlation matrices shown in the topmost panel. GLASSO was fitted using a broad range of tuning parameters, and their relative performance was also compared. GLASSO expectedly performed better than the other competing methods, since it is designed to shrink inverse correlation matrices.

